# Local High-Protein, Plant-Based Ready-to-Use Therapeutic Food Enhances Recovery from Malnutrition in Rats

**DOI:** 10.1101/2024.11.22.624820

**Authors:** Aurélie Bechoff, Peter Akomo, Molly Muleya, Anastasios D. Tsaousis, Charoula Konstantia Nikolaou, Laura Utume, Aviv Schneider, Mona Khalaf, Ram Reifen, Efrat Monsonego-Ornan

## Abstract

Infant child malnutrition is a major public health issue. We conducted a preclinical study with young rats to mimic the conditions of child malnutrition (combined wasting and stunting) and evaluate recovery using a novel plant-based ready-to-use-therapeutic food (RUTF) formulation.

Three-week old female Sprague Dawley rats were assigned to six treatments groups in a 6-week experiment. The treatments included: 1) control balanced diet (CT), 2) A protein-deficient diet to induce malnutrition (MN), 3) and 4) A control balanced diet followed by either commercial RUTF (CT-PM) or a locally produced plant-based RUTF (CT-ChSMS), and 5) and 6) a protein deficient diet followed by either commercial RUTF (MN-PM) or locally produced plant based RUTF (MN-ChSMS), respectively. In treatments 3-6, rats were initially fed either a control-balanced or protein-deficient diet for 3 weeks, followed by 3 weeks of either the commercial or the locally plant-based RUTF.

Results showed that rats in the CT-ChSMS group exhibited growth and weight comparable to CT group, while those in the MN-PM group showed no significant improvement compared to the MN group. Notably, rats in the MN-ChSMS group demonstrated significant catch-up growth, whereas those in the MN-PM group did not.

Additionally, consumption of ChSMS and PM RUTFs differed significantly. ChSMS RUTF which contained 14% protein over total energy with better amino-acid composition and a higher Protein Digestibility-Corrected Amino Acid Score (PDCAAS), resulted in significantly greater weight gain and length compared to PM RUTF, which contained 10% protein over total energy. These findings indicate that a locally produced, culturally acceptable and affordable plant-based RUTF formulated with high protein quality and quantity may be effective in treating acute and chronic malnutrition in children.

## Introduction

Around 45 million children under 5 years old suffer from acute malnutrition, with a third facing the severe form (1) known as severe acute malnutrition (SAM), which contributes to about 1 million deaths annually (2). Acute malnutrition can become chronic if food shortages persist, leading to stunting, which affects 148 million children worldwide (1). Children can also experience a combination of acute and chronic malnutrition, known as underweight. Acute and chronic malnutrition have long-lasting impacts, impairing development, cognition and immune function, increasing the risk of non-communicable diseases in later life (2,3). In poverty-stricken settings, poor hygiene, limited access to clean water and co-existing diseases can exacerbate malnutrition by causing gut inflammation and affecting gut microflora, reducing nutrient absorption (4) and increasing relapse of acute malnutrition (5).

Ready-to-Use-Therapeutic-Food (RUTF) is the current treatment for SAM among children with no medical complications, as prescribed by the World Health Organization (WHO) and can also be used to treat Moderate Acute Malnutrition (MAM) (6). The standard RUTF formulation (PM-RUTF) is typically a paste made with milk powder, peanuts, sugar, and oil and enriched with vitamins and minerals. RUTF was developed 20 years ago, and its nutritional content equates the previous in-hospital formulation (a dried milk powder enriched in vitamins and minerals (F100) developed to meet the nutritional requirements of children aged 6-59 months suffering from SAM). Stunting can be treated using lipid-based nutrient supplements (LNS), which have shown remarkable results recently (7). Their nature (paste-like food) and composition are similar to that of RUTF but they are given in smaller quantities (8). In the future, developing a combined product for acute and chronic malnutrition could be beneficial in addressing this public health burden more economically.

A major constraint of SAM or undernutrition treatment in children is poor access of RUTF in the majority of the low-income countries (LICs) where it is needed. This is mainly because a large proportion of RUTF products are produced in high-income countries as opposed to LICs where they are needed as the main ingredient required in RUTF formulation, skimmed milk powder, is costly and not readily available in LICs. Currently, only 30% of children suffering from SAM are being reached with this treatment (9,10). With the new Codex Guidelines (11), UNICEF and other global institutions have recently enabled the development of alternative non-milk formulations made from locally sourced ingredients, which could stimulate local economies and be more affordable and more environmentally sustainable (12,13). Recently, a novel plant-based RUTF formulated using African-grown ingredients (soybean, maize, and sorghum: SMS) was developed. This formulation improved iron status (14) and restored circulating amino acids in children with SAM (15). Children recovered well, but the weight gain was typically less than with the standard PM-RUTF formulation (i.e., Plumpynut or equivalent (16,17) and this was attributed to the lower protein quality of the SMS

High protein quality is essential for the growth and development of young children, especially those with SAM. In SAM, protein is vital for both maintenance and growth. For weight maintenance, protein typically accounts for about 3% of total energy intake, while protein requirements for catch-up growth in children with SAM are 10-12% of total energy. However, the actual protein needs for optimal growth, considering protein quality, may be higher (18). However, determining the precise amounts and quality of protein required is challenging, as absorption varies depending on food matrix and individual needs, making protein optimisation particularly difficult (19). Moreover, children with SAM require higher protein during infection (20), and those with oedema may experience reduced protein utilization, leading to decreased plasma protein levels (21). The type, and quality of protein could also play an important role in the recovery from acute and chronic malnutrition. A study showed that aromatic amino acids increased protein synthesis in children with SAM (22) and another suggested that milk protein were superior to whey and soya (23). However, other clinical studies did not find difference in MAM children recovery with different protein qualities (24) or types (25).

Animal models are used to effectively replicate biological processes associated with conditions such as malnutrition or disease in humans, providing valuable insight into potential treatments and cures. Preclinical models serve as an effective and cost-efficient intermediate step when testing a new RUTF formulation, offering ethical advantages over clinical trials.

A preclinical study with young Sprague Dawley rats demonstrated varying levels of weight and length gain when different protein sources (casein, soybean, fly, spirulina, chickpea) were incorporated at a level of 10% in the diet (26). The results highlighted that both amino-acid composition and the type of protein played important roles in growth. However, when protein content was increased to 20%, the type of protein had no significant effect on recovery (27,28). These findings suggest that the low protein quality of some ingredients may be overcome by increasing the protein quantity to match that of the higher protein quality sources.

We found only two preclinical studies using Sprague Dawley rats that tested the effectiveness of plant-based RUTF (29,30) but those provided limited insights. In this study, we applied a preclinical model using Sprague Dawley rats, to compare PM-RUTF (10% protein of total energy), with a novel high-protein and plant-based ChSMS formulation (14% protein of total energy) made with SMS and concentrated chickpea. This new formulation is a modification of the SMS formulation previously developed and clinically tested by Akomo and colleagues (14,15).

## Materials and Methods

### In vivo Animal (Rat) Model Experiment

This animal model experiment using female Sprague Dawley (SD) rats was set up to mimic underweight (wasting and stunting) in 6-59-month-old children and then simulate treatment and possible repletion. The 6-week experiment was carried out under standard environmental conditions (23±1°C) with a 12-hour (8am–8pm) light and dark cycle in a Non-SPF (Conventional) animal facility, at the Robert H. Smith Faculty of Agriculture, Food and Environmental Sciences, Hebrew University of Jerusalem Israel (HUJI). All animals were handled according to ethical care and use of laboratory animals, as approved by the HUJI Authority for Biological and Biomedical Models (ABBMS) (Ethics Number AG-22-16962-3). Forty-eight three weeks old recently weaned SD rats were purchased from Harlan Laboratories (Rehovot, Israel). Upon arrival, the rats were weighed and randomly allocated to 12 rats to 4 cages each and individually marked by ear piercing (2 cages or 8 animals per treatment group) (Table 1).

**Table 1:** Experimental treatment groups

| **CODE** | **Treatment Groups** |
| --- | --- |
| CT | Standard rodents diet for growth (control group) |
| MN | Protein deficient-diet inducing malnutrition (malnourished group) |
| CT-PM | CT group switched to commercially available RUTF |
| MN-PM | MN group switched to commercially available RUTF |
| CT-ChSMS | CT group switched to ChSMS plant-based RUTF |
| MN-ChSMS | MN group switched to ChSMS plant-based RUTF |

Prior to the start of the experiment, rats were given a 3-day acclimation period, during which they were fed regular rat chow and water.

A 6-week-long experiment was conducted on SD rats after weaning. This time frame (from 3-9 weeks old) was selected to mimic the human growth period up to sexual maturity (31). During this accelerated growth period, the rats were initially fed the recommended diet for growing rodents (CT) or a protein and micronutrient-deficient diet (MN), simulating underweight, for 3 weeks. After 3 weeks, the rats from both diets received either the tested novel ChSMS formula (CT-ChSMS and MN-ChSMS), the commercial PM formula (CT-PM and MN-PM), or continued with the same diets (CT and MN) for more 3 more weeks (Figure 1).

**Figure 1:**
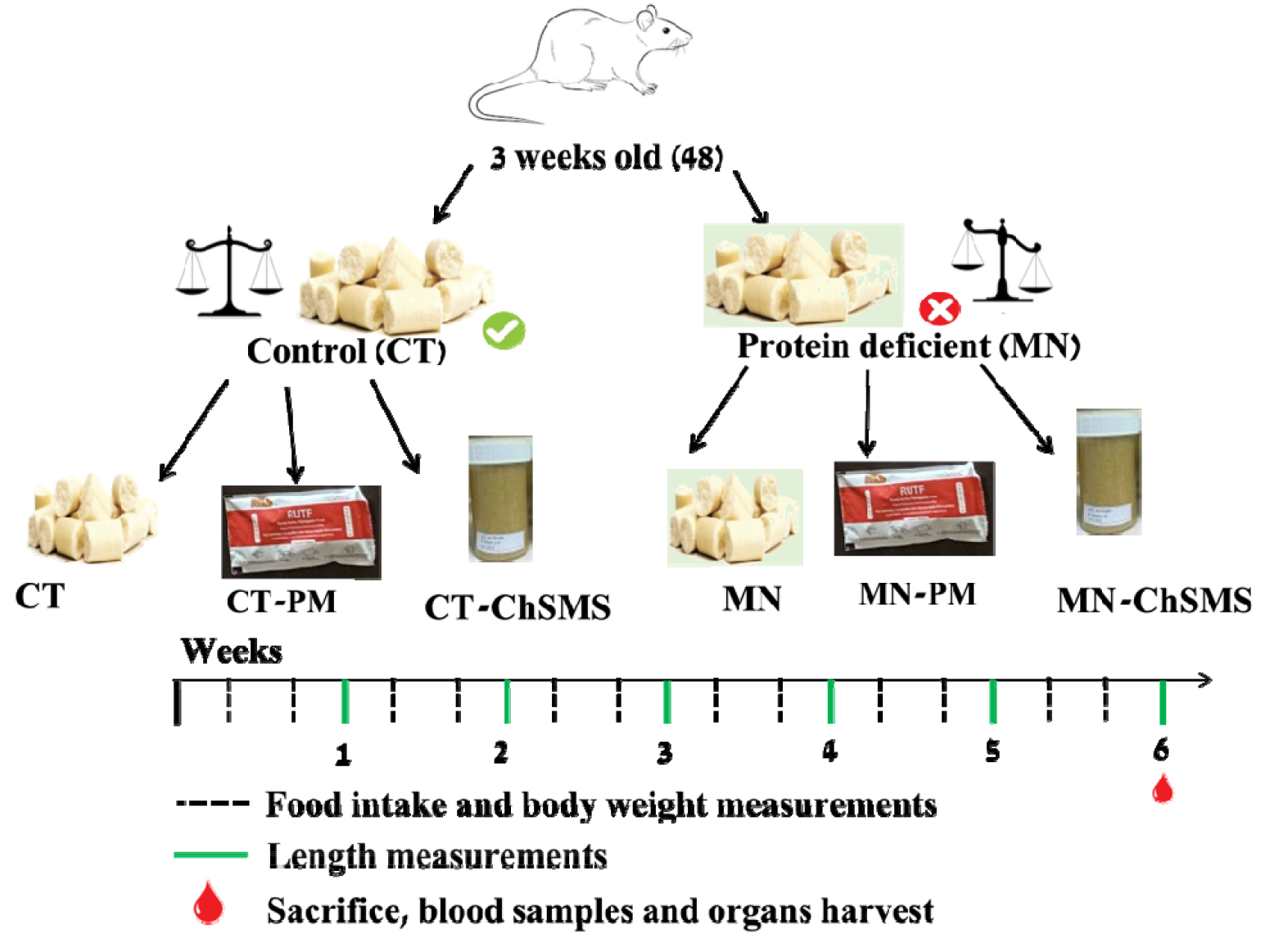
Schematic Diagram of Preclinical Experimental Set-up

Throughout the experiment period, food consumption and body weight were recorded twice a week, while body length (measured from the tip of the nose to the end of the tail) was measured once a week. Additionally, rats were monitored for any physical and behavioural changes (Figure 1).

### Diet Composition and Preparation

The rats were given four (4) diets throughout the duration of experiment. Three (3) diets were prepared in the laboratory under sanitary conditions and one (1) purchased from commercial producers and suppliers.

- **Control balanced diet (CT)**: this was formulated at HUJI based on the American Institute of Nutrition (AIN-93G) recommendation for the growth phase of rodents – 16 % fat, 63.5 % carbohydrates and 20.5 % protein; with the addition of multimineral and multivitamins based on the same recommendation (Table 2).

**Table 2:** Ingredient composition of balanced and unbalanced experimental diets (values are presented as grams per kg dry weight)
- **Unbalanced Protein-deficient diet (MN)**: this was also formulated at HUJI based on the AIN-93G recommendation for unbalanced diet – 25 % fat, 70 % carbohydrates and 5 % protein; the addition of only 50 % of multimineral and multivitamins recommended (Table 2).

| Product/Ingredient | Control balanced diet (CT)<br>(g/kg diet) | Unbalanced diet (g/kg diet) (MN) |
| --- | --- | --- |
| Cornstarch | 397 | 438.7 |
| Dextrinized Cornstarch | 132 | 145.7 |
| Sucrose | 100 | 110 |
| Fibre | 50 | 50 |
| Soybean Oil | 70 | 120 |
| Casein ($\geq 85\%$ Protein) | 200 | 50 |
| L-Cystine | 3 | 3 |
| Mineral mix (AIN-93G-MX) | 35 | 17.5 |
| Vitamin mix (AIN-93G-VX) | 10 | 5 |
| Choline bitartrate (41.1% Choline) | 2.5 | 1.34 |
| Tert-Butylhydroquinone | 0.014 | 0.015 |

Commercially purchased ingredients based on the AIN-93G recommendation were separately weighed and added to a large plastic tub. The ingredients were weighed and thoroughly homogenized and adjusted with water to form a firm dough. The dough was then moulded into medium-sized dumpling patties and air-dried in an extractor hood chamber for 42 hours at ambient temperature. Air-dried dumpling patties were weighed, packaged into portions of approximately 250 g, appropriately and stored at -20 °C until use.

#### Chickpea, Sorghum, Maize, Soybeans (ChSMS) RUTF

initially, several RUTF formulations were theoretically developed from linear programming (32) to meet the 2022-UNICEF RUTF Specifications and Codex requirements. Then, a prototype production (2.5kg per formulation) was conducted under sanitary conditions in the laboratory at Equatorial Nuts Processors (ENP), Murang’a, Kenya in May and November 2022. Samples of the prototypes were shipped and tested for nutritional conformance at Eurofins Laboratories, UK. Following this, ChSMS was selected as the most suitable formulation and prepared in larger quantities at ENP (20kg). Its nutritional composition was re-analysed at Eurofins Laboratories, checked for aflatoxin level and microbiological safety at ENP, and its amino acid measured at the Nutritional Composition and Digestibility Lab (NCDL) of the University of Nottingham, UK.

#### Peanut butter and milk (PM) Standard RUTF

the commercially available standard RUTF was purchased from Nuflower Foods and Nutrition Private Limited, Haryana, India. Its nutritional composition was verified at Eurofins UK.

ChSMS and PM were shipped to HUJI, Rehovot, Israel for the experiment (Table 3).

**Table 3:**
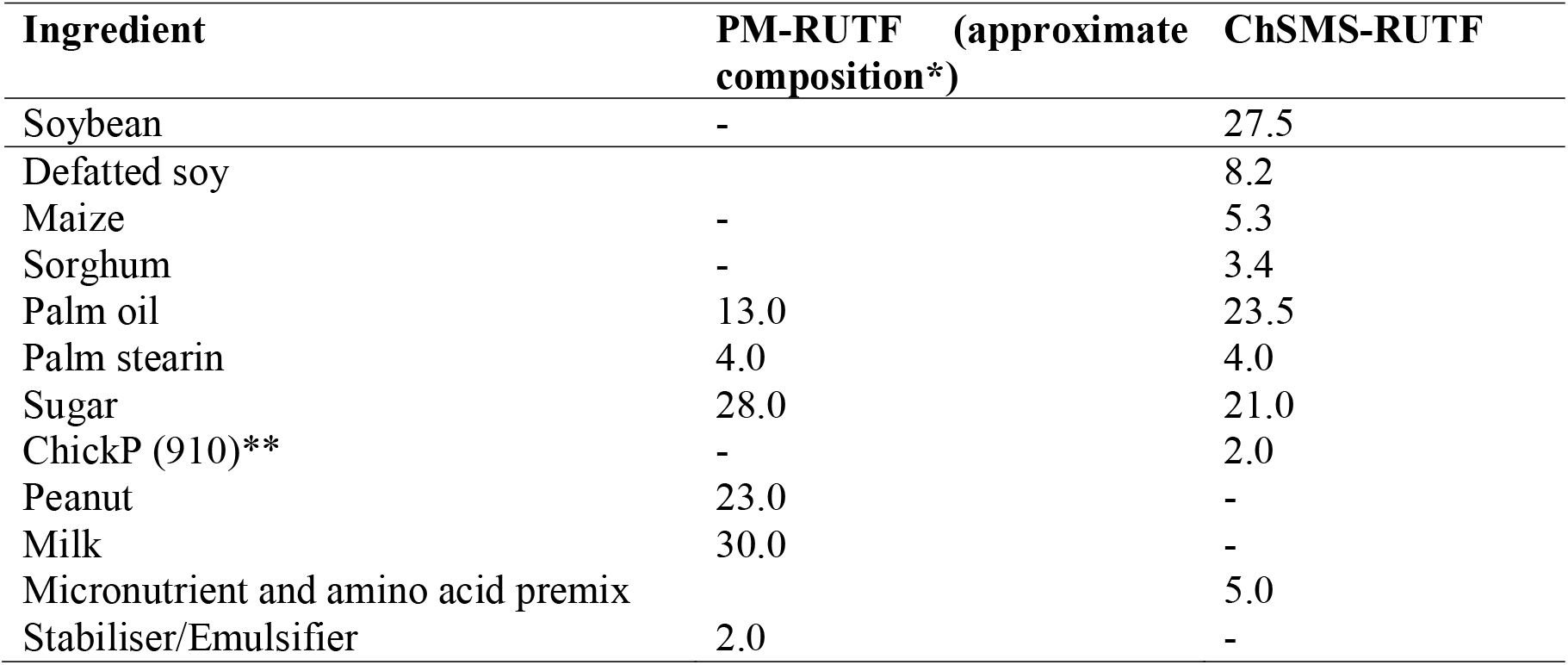
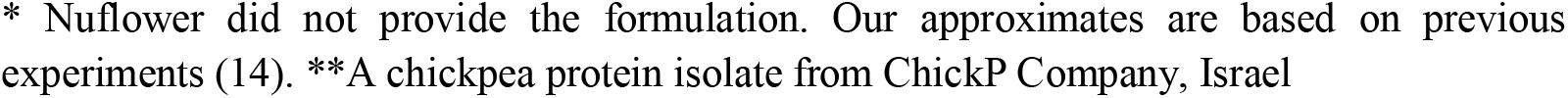
Ingredient composition of ChSMS and PM RUTFs (values are presented as percentage weight)

| Ingredient | PM-RUTF (approximate composition*) | ChSMS-RUTF |
| --- | --- | --- |
| Soybean | - | 27.5 |
| Defatted soy |  | 8.2 |
| Maize | - | 5.3 |
| Sorghum | - | 3.4 |
| Palm oil | 13.0 | 23.5 |
| Palm stearin | 4.0 | 4.0 |
| Sugar | 28.0 | 21.0 |
| ChickP (910)** | - | 2.0 |
| Peanut | 23.0 | - |
| Milk | 30.0 | - |
| Micronutrient and amino acid premix |  | 5.0 |
| Stabiliser/Emulsifier | 2.0 | - |

\* Nuflower did not provide the formulation. Our approximates are based on previous experiments (14). \*\*A chickpea protein isolate from ChickP Company, Israel

## Results

### Nutritional composition of the commercial versus high-protein plant based ready-to-use Therapeutic Foods used for the preclinical study

Nutritional composition data of the commercial PM-RUTF and the novel ChSMS-RUTF are described in Table 4.

**Table 4:**
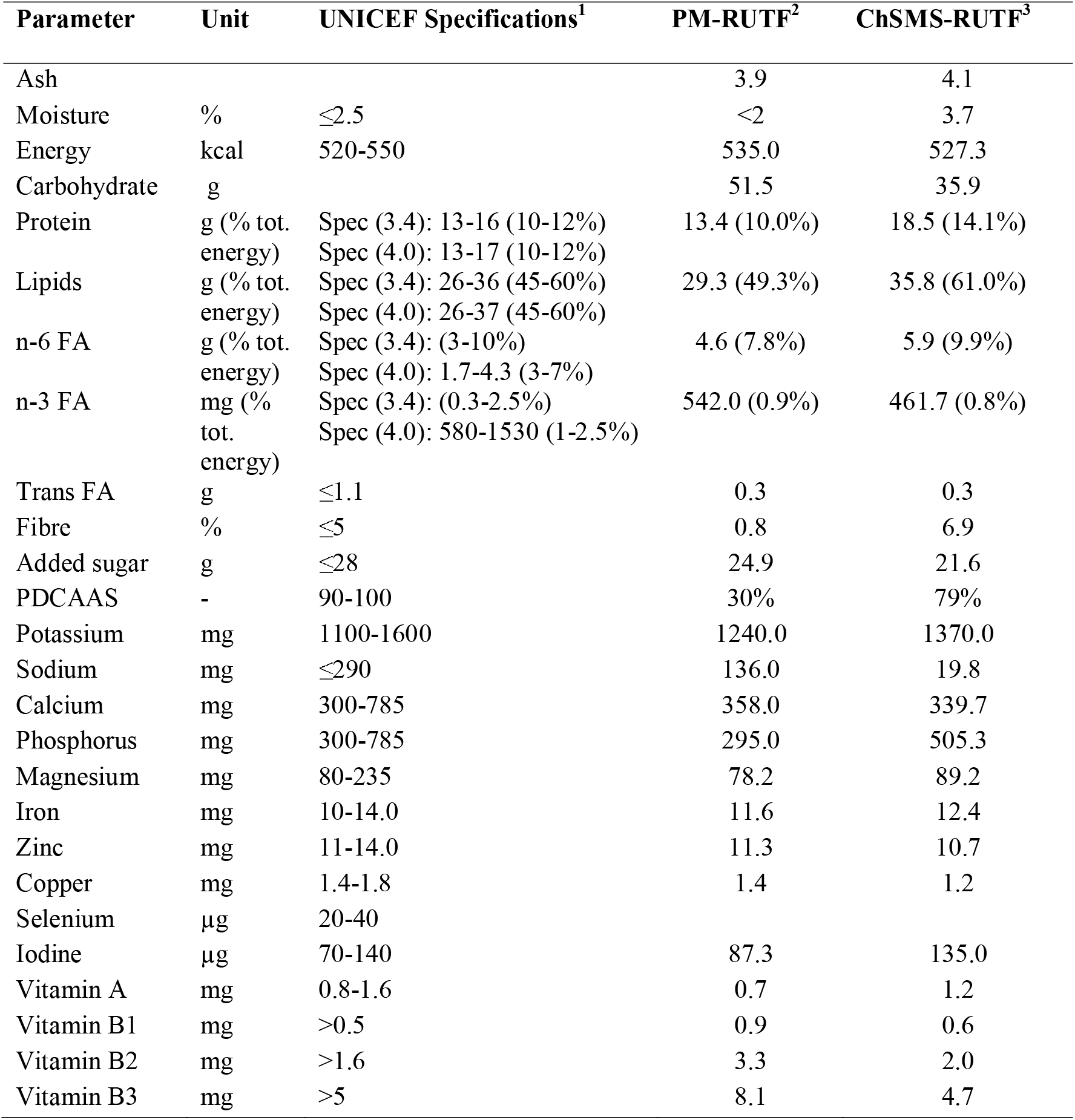
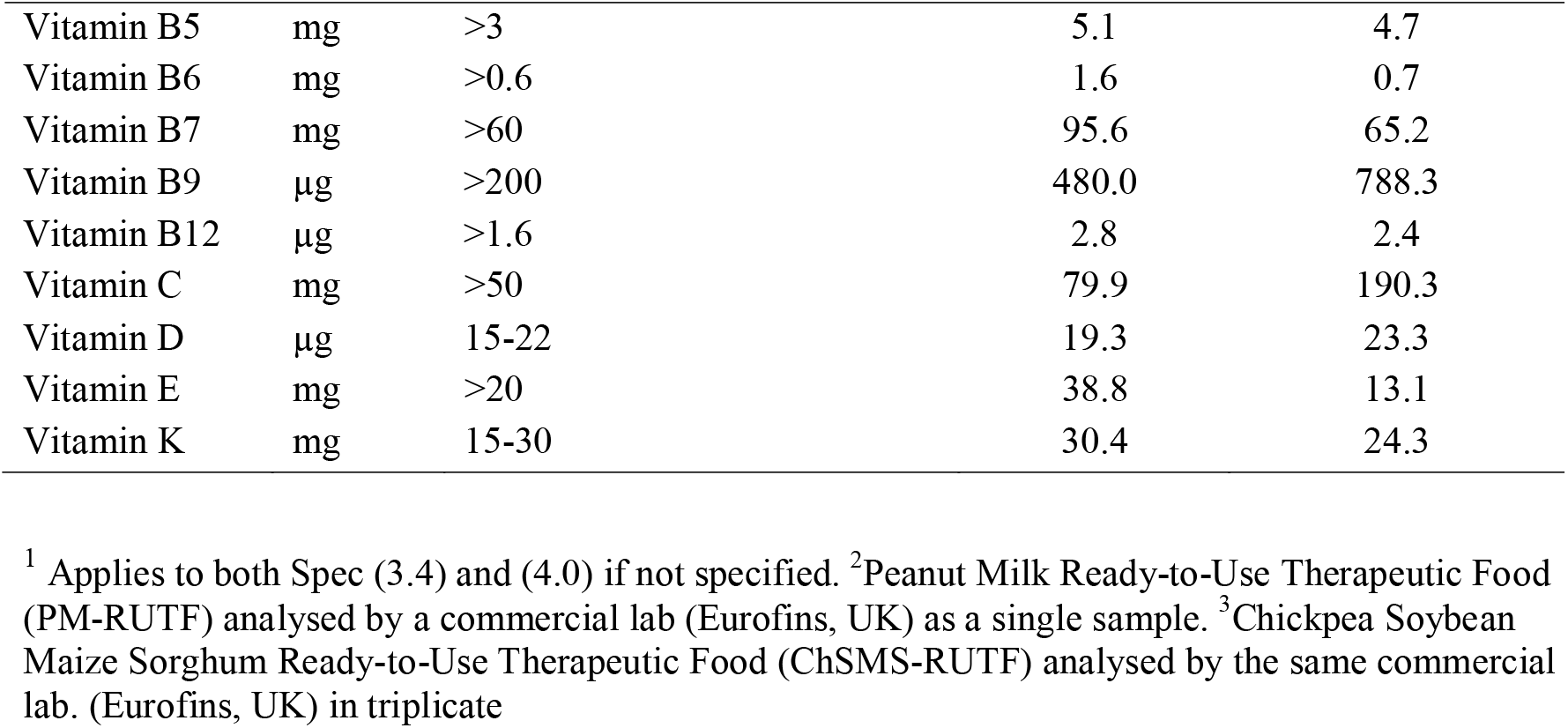
PM-RUTF and ChSMS RUTF nutritional composition (per 100 g)

### The Effect of the new formula on growth performance in a rat model for severe malnutrition and stunting

The experimental period was divided into two phases: the first 3 weeks generated control (normal) (CT) and malnutrition (underweight) (MN) status whilst the last 3 weeks feeding on either PM-RUTF or ChSMS-RUTF simulated RUTF treatment and potential recovery.

The effect of the diets on growth was first determined in terms of body weight, representing body growth including energy balance, and length from tip to nose as well as lengths of the femora as the longitudinal bone growth parameters (Fig 2). Body weight (BW) comparison between the groups demonstrates the effect of the deficient diet (MN), that was clearly detected already after 4 days (Fig 2A, B). BW differed significantly between the groups and continued to increase, such that after 3 weeks, BW of the rats were 167±20g and 79± 6g for CT and MN groups respectively. The same pattern was recorded for rats’ length (Fig 2C, D), with significant length differences after 10 days, and total body length of 35+1cm and 29+1cm for CT and MN groups respectively, after 3 weeks of feeding. These results demonstrate malnutrition and stunting in the group fed with deficient diet.

**Figure 2:**
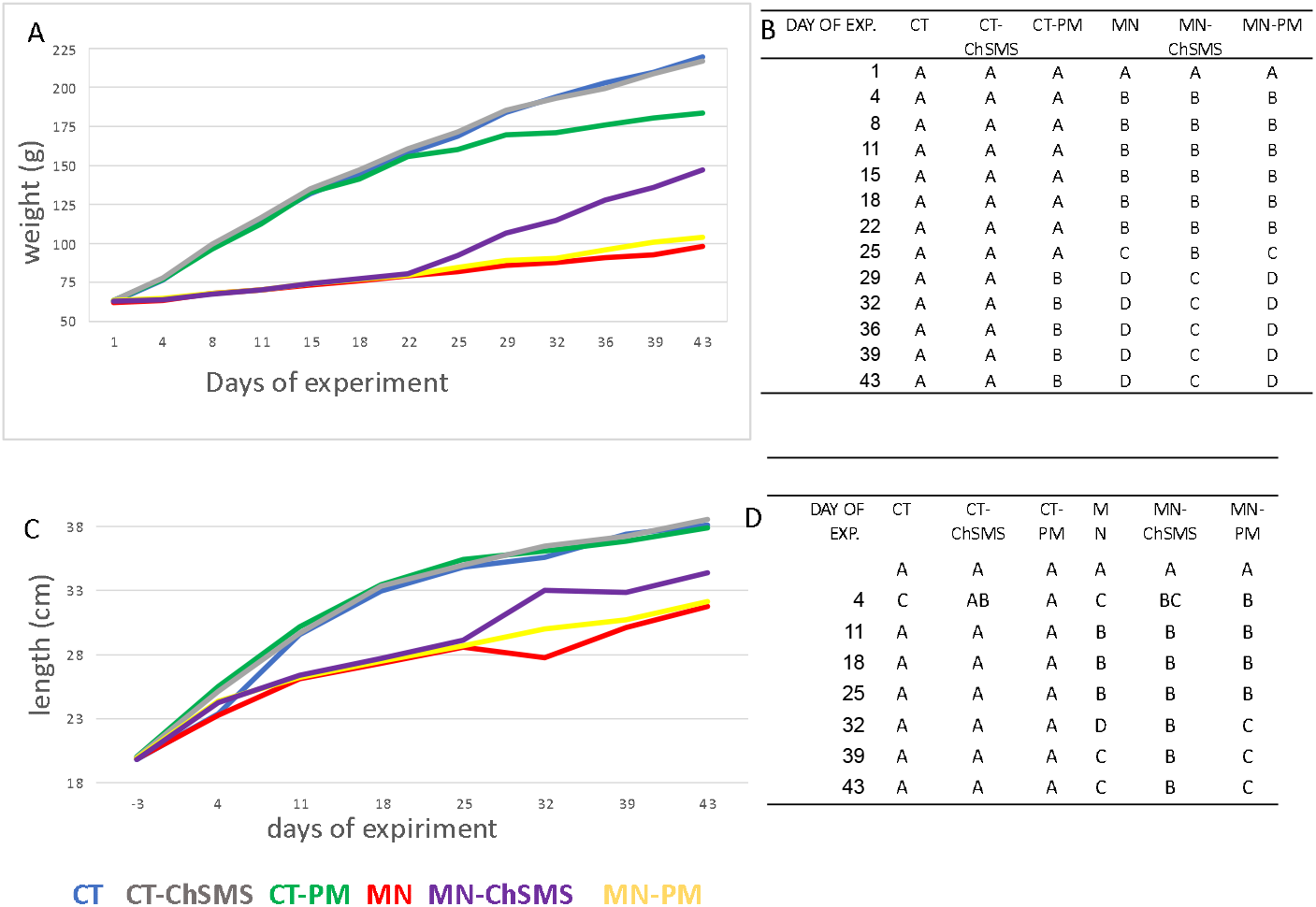
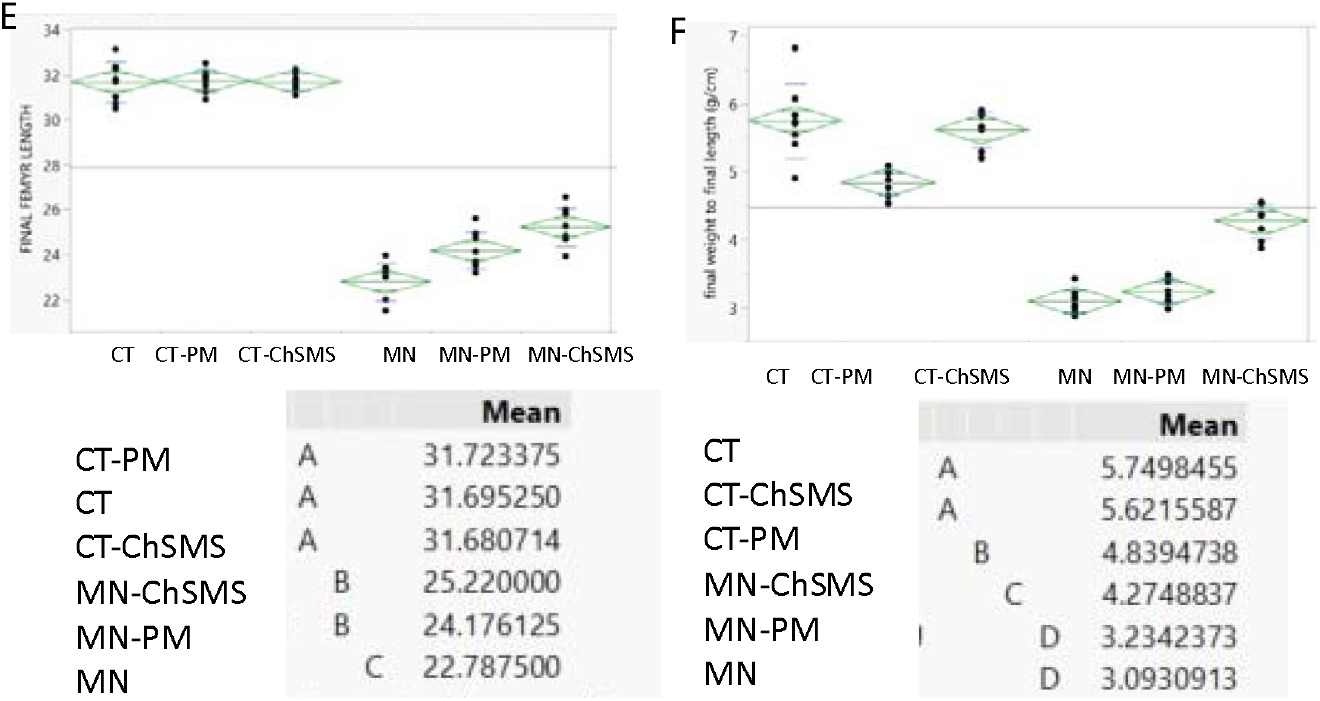
Growth pattern, (A) Weight (B) BW statistical analyses (C) length, along the exp (D) length statistical analyses (E) final femur length (F) final weight/length

After 3 weeks, the diets were replaced with RUTF formulas in order to achieve rescue and catchup growth. Surprisingly, the commercial formula (PM) did not lead to increased growth (BW and length) in the malnourished animals (MN-PM) (Fig 2A-D). On the other hand, the malnourished rats fed with the novel formula (MN-ChSMS) improved BW after 4 days of diet shift and longitudinal growth 10 days post initiation of RUTF consumption (Fig 2A-D). These trends continued until the end of the experiment, at which rats fed with the ChSMS formula reached a BW of 147g and a length of 34cm, which were significantly higher than those of the malnourished rats (98g and 32cm). However, the initially malnourished rats did not achieve full recovery, and did not reach the same BW and length of control rats (220g and 38cm) along the whole period. In contrast, rats fed on the PM formula did not differ from those consuming the deficient diet (MN vs. MN-PM) (Fig 2A-D). When replacing the control diet (CT), the ChSMS formula did not affect BW nor length, but the PM one, led to slower growth in BW without an effect on body length (Fig 2A-D).

Femora length, of all rats were measured following sacrifice after 6 weeks of the experiment. The results are the direct measurement of long bone, approximating the data from whole body length measurements (Fig 2E). The longest averaged femur (31.7mm) was in the 3 groups consuming the control diet in first 3 weeks without any effects of the formulas (CT, CT-PM, CT-ChSMS). The shortest femur was in the MN group (22.8mm), then the MN-PM group (24.2mm), and the MN-ChSMS performed the best, but did not achieve a complete catch-up in bone elongation (25.2 mm). To simulate weight for length (W/L) parameter in the rats, final weight was divided by the final length at the termination of experiment. The data show that the ChSMS significantly improved W/L in the malnourished animals and did not affect it in the well-nourished animals (Fig 2F). Altogether the results demonstrate the better ability of the new formula in improving the growth of malnourished (underweight) rats.

### Food consumption of ChSMS and PM formula in a rat model for severe malnutrition and stunting

Throughout the experiment, food intake was measured for calculating energy and macronutrients consumption, to evaluate their contribution to growth (Fig 3). During the first 3 experimental weeks, the rats consumed a higher amount (g) of the CT diet as compared to the MN diet (Fig 3A). Despite the MN diet having higher caloric density compared to CT (3.2 vs 3.5Kcal/g respectively), the MN group also consumed less energy (Kcal) during this time (Fig 3B). This is further highlighted when comparing the total amount and energy consumption of each rat along the whole 6 weeks experiment (Fig 4A,C).

**Figure 3:**
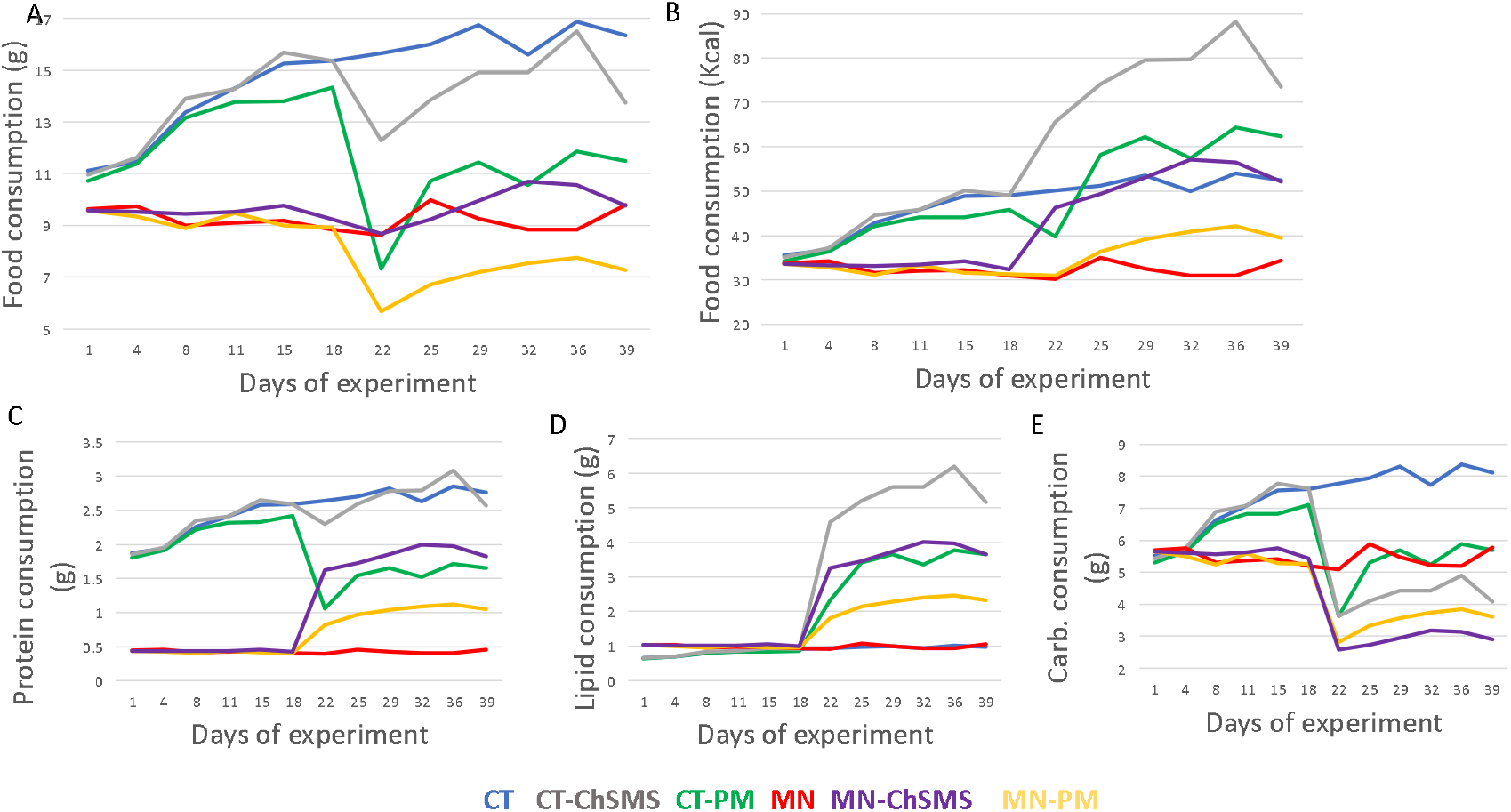
Food Consumption (A) 0-6 weeks (g) (B) 0-6 weeks (kcal) (C) 0-6 weeks protein (g) (D) 0-6 weeks lipids (g) (E) 0-6 weeks carbs (g)

**Figure 4:**
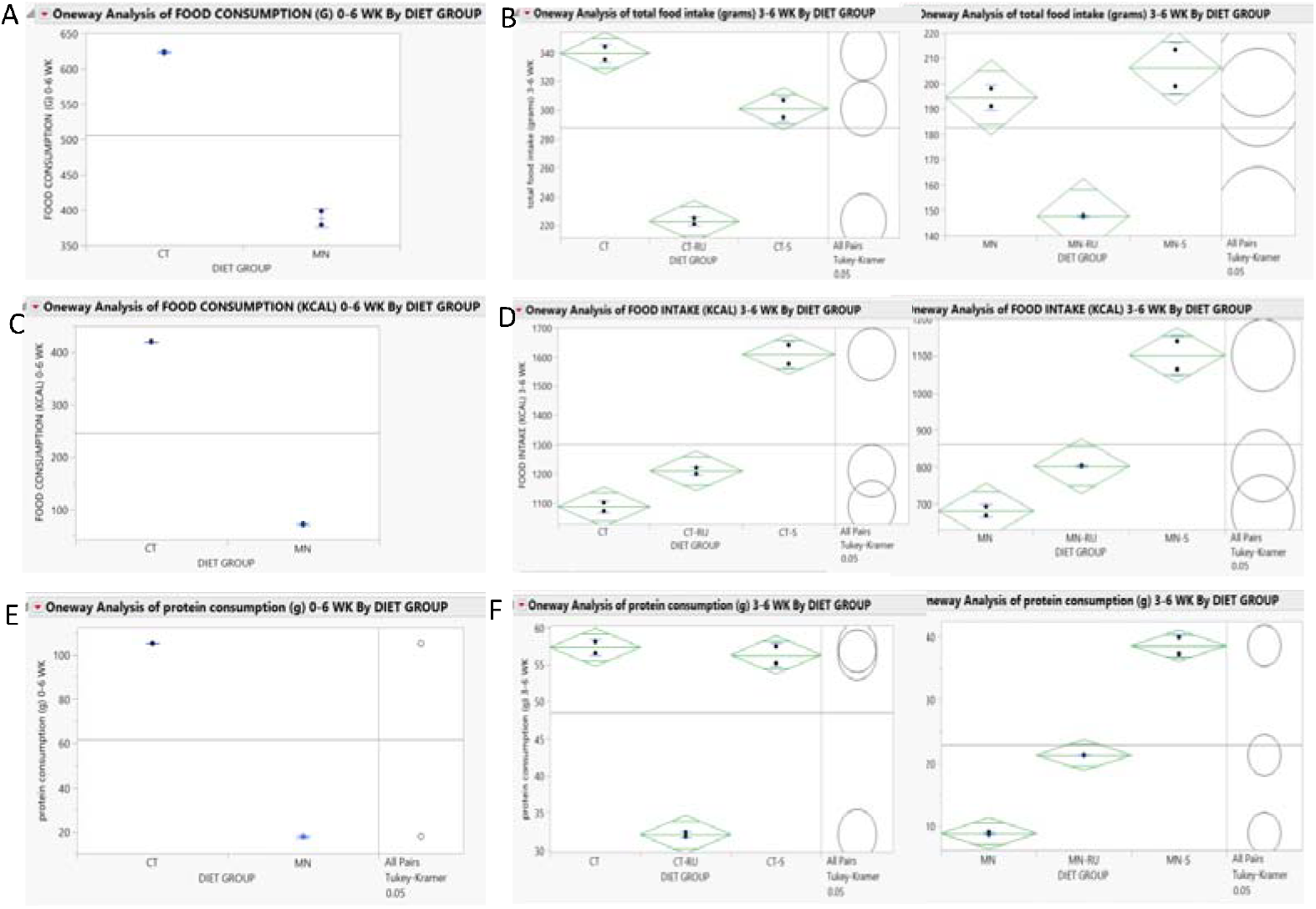
Energy and Protein consumption (A) 0-3 weeks food CT vs. MN (g) (B) 3-6 weeks food (g) (C) 0-3 weeks CT vs. MN (Kcal) (D) 3-6 weeks (Kcal) (E) 0-3 weeks protein CT vs. MN (g) (F) 3-6 weeks protein (g)

Figures 3C and 3D demonstrate the clear benefits of the PM and ChSMS formulas over the MN diet in terms of protein and lipid consumption. Their higher protein and fat content explain it, as carbohydrates constitute 36% of the formulas were consumed in significantly less quantities (Figure 3E). The disparity in total protein consumption between rats on the CT diet and rats in the MN group becomes apparent when examining the data from the 6-week experiment (Fig 4E). A rat on the CT diet consumed 5 times the amount of protein compared to a rat in the MN group.

Energy and protein intake in the CT and MN groups were consistent with their growth performance. However, while in the CT group rats increased their food consumption and grew (BW-from 63g to 220g, and length from 20cm to 38cm), the rats consuming the deficient diet did not increase their food consumption, despite a small increase in BW (from 62g to 98g) and length (from 20cm to 32cm) (Fig 3A,B and Fig 2A,B).

After switching to the formulas, a reduction in food intake (in grams) was recorded for the PM formula in all rats, and for the ChSMS formula in the well-nourished rats but not in the malnourished rats (Fig 3A, 4B). Energy intake revealed an increase for the ChSMS formula for all rats, and no change in the PM, as compared to the CT and MN diets (Fig 3B). This resulted from the higher caloric density of the formulas, i.e., 5.35 kcal/g and 5.27 kcal/g for the PM and the ChSMS formulas, as compared to 3.2 and 3.5Kcal/g for the CT and MN diets, respectively (Table 4).

When comparing the total amount of energy (0-6 weeks) and that consumed between 3-6 weeks by each rat in the experiment (Fig 4C,D), the effects of the formulas are emphasised as compared with the basic diets (CT and MN). Rats that were malnourished (MN) increased their energy consumption during the last 3 weeks from 682 kcal in the MN diet to 803 kcal with the PM and 1102 kcal with the ChSMS formula (Fig 4D). The well-nourished rats (CT) also increased their energy consumption (Fig 4D).

The MN group had lower protein consumption compared to CT, attributed to a lower amount in their diet and reduced food intake in this group (Fig 4E). Both RUTF formulas led to significant increase in protein consumption in the malnourished rats (Fig 3C, 4F), due to their higher protein content of 13.4% and 18.5 % for the PM and the ChSMS formulas, respectively, as compared to 5% in the MN diet (Table 4). While the PM formula led to a 2.4-fold increase, the ChSMS led to a 4.3 times-fold increase than MN. These increased energy and protein consumption by switching from MN diet to ChSMS formula explain the significant increase in BW and length of the whole body and femora (Fig 2).

### ChSMS and PM formula Utilization for Growth

In an attempt to determine dissimilarities in formula quality, we calculated the energy and protein efficiency ratio (EER and PER). EER was calculated as total weight gain per 100 kcal intake and PER measured the body weight gain per gram of consumed protein (Fig 5). We first compare the CT and MN diets in the first 3 weeks of the experiment. Fig 5 A,Cillustrates the comparison between the CT and MN diets during the initial 3 weeks of the experiment. The CT diet showed a higher EER, indicating the inadequacy of the MN diet for supporting healthy growth. Nevertheless, PER remained consistent across both diets, suggesting that protein levels were the limiting factor for growth performance at this stage. The comparison of diets and formulas in the second part of experiment (weeks 3-6) shows that the ChSMS formula was superior in supporting catch up growth in malnourished rats, both in terms of energy and protein (Fig 5 B,D).

**Figure 5:**
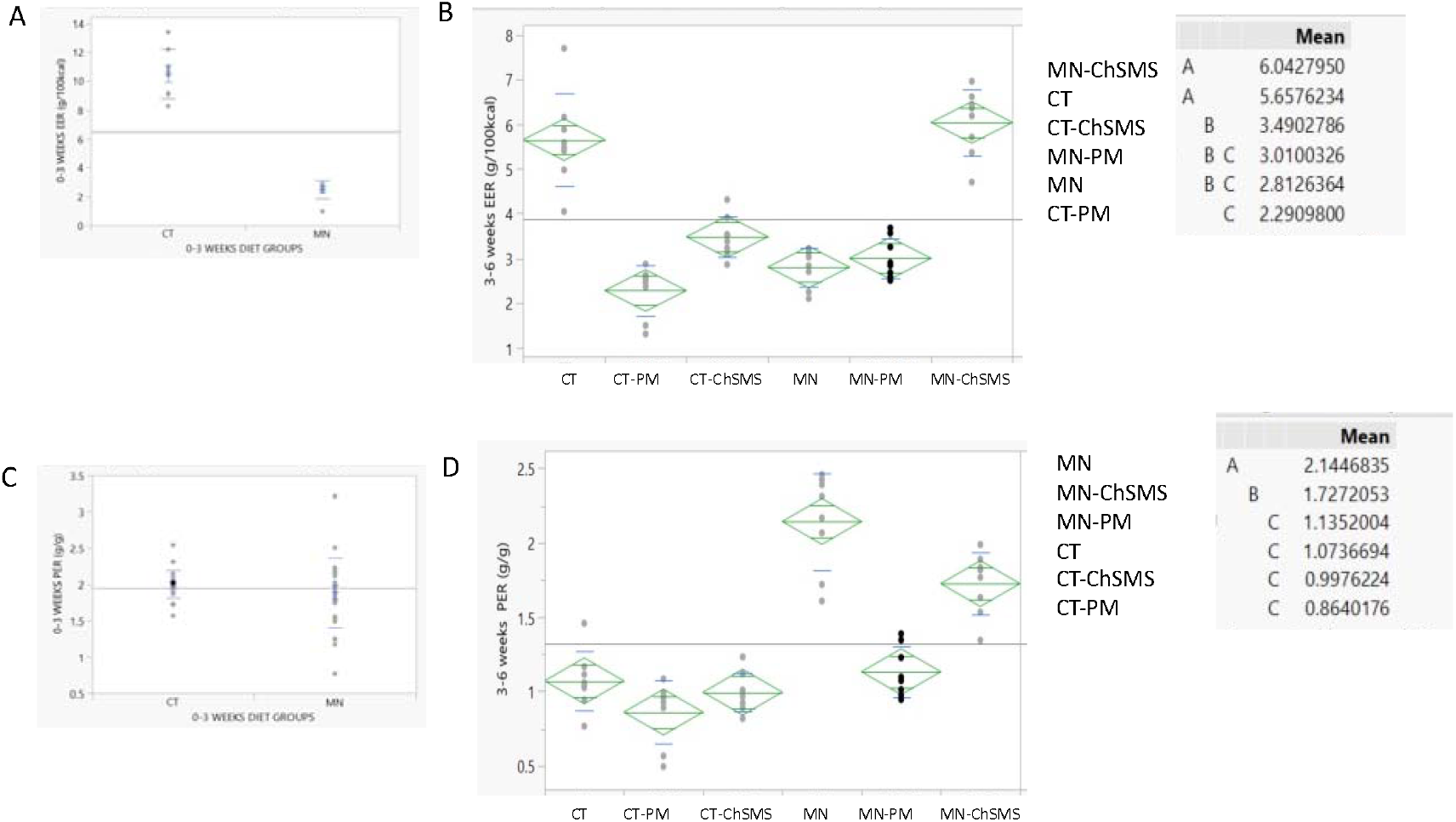
EER and PER (A) 0-3 weeks EER (g/Kcal) (B) 3-6 weeks EER (g/Kcal) (C) 0-3 weeks PER (g/g) (D) 3-6 weeks PER (g/g)

## Acknowledgments

This research was supported by UK Research and Innovation (UKRI) Global Challenges Research Fund (GCRF) 2022-23, Universities UK International (UUKi) UK-Israel Mobility Scheme call 1 2023-24, and Research Excellence Framework (REF) funds from the University of Greenwich (2021-23). We are grateful to Dr. Moses Mwangi for allowing us the use of the Equatorial Nuts Processors (ENP) laboratories to carry out the formulation of our novel RUTF. We thank Samuel Maina for providing the assistance of laboratory technicians to support the formulation process, Emma Mugo (head of the laboratory), Laurine, Lucky, Charles, Brian, Mary, with the participation of Benard Maguti the Quality Assurance Manager. We are also grateful to Sveta Penn, the laboratory technician and students, Jerome Janssen, Aviv Schneider, Mona Khalaf who helped support the animal study at HUJI. The opinions expressed in this paper do not necessarily reflect the views of our donor or partners.

